# Chronic broadband noise increases the fitness of a laboratory-raised freshwater zooplankton

**DOI:** 10.1101/2022.11.19.517212

**Authors:** Loïc Prosnier, Emilie Rojas, Olivier Valéro, Vincent Médoc

**Affiliations:** ENES Bioacoustics Research Team, CRNL, University of Saint Etienne, CNRS, Inserm, Saint-Etienne, France; France Travail, Saint-Etienne, France; Department of Biology, Norwegian University of Science and Technology (NTNU), Trondheim, Norway

**Author notes:** **Corresponding author:** Loïc Prosnier.

**Keywords:** *Daphnia magna*, Acoustic pollution, Broadband noise, Fitness, Mobility

## Abstract

Although there is an increasing interest in the effects of anthropogenic noise on underwater wildlife, most studies focus on marine mammals and fishes while many other taxa of substantial ecological importance are still overlooked. This is the case for zooplankton species which ensure the coupling between primary producers and fishes in pelagic food webs. Here, we measured lifespan, reproduction, and mobility of laboratory-raised water fleas *Daphnia magna*, a widespread freshwater zooplankton species, in response to continuous broadband noise. Surprisingly, we found a significant increase in survival and fecundity, leading to a higher individual fitness when considering total offspring production and a slight increase in the population growth rate according to the Euler-Lotka equation. Exposed water fleas were found to be slower than control individuals, and we discussed potential links between mobility and fitness. Our results can have implications in aquaculture and for in-lab studies (e.g., in ecotoxicology) where the acoustic environment receives little attention. Chronic broadband noise can be associated with certain human activities, but the consequences for natural *Daphnia* populations might differ as reduced velocity could have negative outcomes when considering competition and predation. Our work is one of the few showing an effect of noise on individual fitness and suggests that noise should be better accounted for in laboratory studies.

## Introduction

Underwater sounds can be of biological (i.e., vocalisations), geological (i.e., rain and wind) and anthropogenic (i.e., human activities) sources and serve as important cues for reproduction, orientation, settlement, territoriality and foraging in both vertebrates and invertebrates (Popper et al., 2020). Anthropogenic noise is now recognised as a ubiquitous pollutant (Hildebrand, 2009; Xie et al., 2011; Frisk, 2012; Rountree et al., 2020) as well as light or chemicals. The noise emissions from seismic surveys, sonar, or pile-driving activities are short and intense (“peak”), while those from recreational and commercial shipping or wind farms are less intense but last longer (“continuous” and “intermittent”) (Francis & Barber, 2013). The frequency range of anthropogenic noise (<2 kHz) overlaps with that used by most animals (Duarte et al., 2021) and many studies document the negative effects on terrestrial and aquatic wildlife (Shannon et al., 2016; Popper & Hawkins, 2019), with greater attention to the marine environment compared to fresh waters (Williams et al., 2015; Sole et al., 2023 ; but see Mickle & Higgs, 2018). Examples of noise-induced alterations in behaviour and physiology abound in the literature (Richardson et al., 1985; Wysocki et al., 2006; Read et al., 2014; Radford et al., 2014; Nedelec et al., 2017; Rojas et al., 2021; Tremblay et al., 2025) but consequences beyond the individual level, for instance on population and community dynamics, remain to be explored (Reid et al., 2019; Rojas, Gouret, et al., 2023).

Another crucial step toward understanding the ecological impact of noise is to investigate the response of invertebrates (Jerem & Mathews, 2021; Sole et al., 2023; Davies et al., 2024). Despite calls for increased research on invertebrates (Morley et al., 2014) and a growing number of studies focusing on crustaceans (Celi et al., 2013), molluscs (Hubert et al., 2022), cnidarians (Sole et al., 2016), and insects (Villalobos-Jime nez et al., 2017), many invertebrates remain underexplored. This is the case for zooplankton (Vereide & Ku hn, 2023; Prosnier, 2024; Davies et al., 2024), which includes small protozoans to large metazoans with gelatinous species (jellyfish, scalps and ctenophores), rotifers, crustaceans but also larvae of fishes, molluscs, bryozoans, and insects. Zooplankton plays a pivotal role in the functioning of pelagic food webs (Ratnarajah et al., 2023) coupling primary producers with higher trophic levels consumers, and is commonly used as bioindicators in ecological quality assessments (Jeppesen et al., 2011; Mun oz-Colmenares et al., 2021). Although zooplankton species generally lacks specialized hearing systems (but see the chordotonal organ in some crustaceans), they possess external mechanoreceptors capable of detecting changes in particle motion (Popper et al., 2001; Nedelec et al., 2016; Sole et al., 2023). For instance, copepods have such receptors on their antennae (Gassie et al., 1993; Weatherby & Lenz, 2000) and have shown behavioural changes to emitted vibrations in water (Buskey et al., 2002) and to sound (Waggett & Buskey, 2008; Tremblay et al., 2025). Only one study showed mechanoreceptors for cladocerans, in their distal swimming and filtering setae (Klann & Stollewerk, 2017). Environmental sound is an important cue for zooplankton species because it enables them to orient and to detect prey and predators (Yen & Okubo, 2002; Chan et al., 2010; Buskey et al., 2011).

Evidence suggests that the effects of noise pollution on zooplankton differ depending on the species and sound features (frequence and frequency variations). Short-term exposures (less than 4 days) to air gun (an intense acute source, 150-180 dB SEL Re 1 µPa^2^) have been shown to cause death of several marine species (McCauley et al., 2017) and negatively impact copepod physiology and development (Fields et al., 2019; Vereide et al., 2023). Conversely, short exposures to less intense motorboat noise (i.e., few days at 126 dB RMS Re 1 µPa^2^) seem to not alter the behaviour of the water flea *Daphnia magna* (Sabet et al., 2015, 2019; Rojas, Prosnier, et al., 2023) but have been shown to reduce the feeding rate of copepods (Ku hn et al., 2023) and decrease the fecundity of rotifers (Aspirault et al., 2023). For short-lived organisms, chronic noise exposure can easily last a significant portion of their lifespan or even exceed it (Barber et al., 2010; Kok et al., 2023). The noise exposure (+ 20 dB at 110-120 Hz) during all juvenile development (i.e., 9-11 days) of a calanoid copepod, affected development duration as well as adult behaviour (Tremblay et al., 2025). Thus, when studying zooplankton, it is important to consider the consequences on fitness (defined as lifetime reproductive success) as the absence of behavioural response does not preclude detrimental effects on survival and/or fecundity. In a previous study, we exposed the water flea *Daphnia magna*, a common freshwater zooplankton model used in ecotoxicological studies (Lampert, 2011; Reynolds, 2011; Bownik, 2017, 2020; Tkaczyk et al., 2021; Ebert, 2022), to playback of nautical activity from birth to death and found no effect on survival or fecundity (Prosnier, Rojas, et al., 2023). To date, only two long-term studies have investigated zooplanktonic communities exposed to boat noise, with or without fish as top-predators (Rojas, Desjonque res, et al., 2023; Rojas, Gouret, et al., 2023). These studies revealed complex outcomes, with different impacts across species, for instance with a positive effect on Bosminidae but negative effects on Daphniidae.

When considering successive boat passages as a sequence of acute stimuli, even if the individual acoustic signatures are not exactly the same, nautical activity can be classified as a form of chronic intermittent stress (Ladewig, 2000). If the impact of noise is dose-dependent (Tyack & Thomas, 2019), for instance through the simulation of mechanoreceptors, then a continuous noise exposure compared to what organisms could experience near water catchment systems or in aquatic husbandries may exert a greater impact than nautical activity (intermittent noise (Prosnier, Rojas, et al., 2023; Vereide et al., 2023)) and short-term acute exposure, depending to the total amount of noise (Fields et al., 2019; Ku hn et al., 2023). To test this hypothesis, we conducted a laboratory experiment where isolated *D. magna* were exposed to continuous broadband noise throughout their entire life (approximately one month, from birth to death). We measured key fecundity parameters including clutch size, clutch frequency, total offspring production and survival, allowing direct comparison of results with previous studies (Prosnier, Rojas, et al., 2023), but also morphological and behavioural traits with body size and swimming speed, which are rarely taken into account in studies on the chronic effects of noise (Prosnier, 2024). Assuming a dose-response relationship, we expected continuous broadband noise to be more stressful than chronic intermittent noise such as that associated with boat traffic, which could lead to decreased fitness in *D. magna*.

## Material and Methods

### Animal collection and maintenance

*Daphnia magna* were purchased from Aqualiment (Grand Est, France) 2 months before starting the experiment. They were stored in two 20-L aquariums (50 *D. magna*/L), filled with aged tap water (see on Zenodo repository for the physicochemical composition (Prosnier et al., 2022)) at 18°C, and under a 12:12 light:dark cycle. Every two days, *D. magna* were fed with 0.05g powder of algae, a mix of 80% of *Arthrospira platensis* and 20% of *Aphanizomenon flos-aquae* (Algo’nergy® Spiruline + Klamath, from organic agriculture), *per* aquarium. The use of algal powder both in the rearing tanks and the experimental microcosms limits the risk of bias due to live food, whose temporal and spatial dynamics could be influenced by noise (it seems unlikely to us that noise affects algae sedimentation).

### Fecundity and mortality

The protocol is similar to that of Prosnier, Rojas, et al. (2023), designed to perform experiments on isolated individuals from birth to death (Fig. 1a). Gravid *D. magna* were collected (from the storage aquarium) and isolated in 50-mL jars containing Volvic® water, known as a good medium for *D. magna* (B. Prosnier from Eau de Paris, a French water industry, pers. com.). Newborns (<24h) were individually transferred into 150-mL (5.6 x 8.6 cm) glass-microcosms (one identified individual *per* microcosm to track their fecundity and mortality), sealed at the top with a 0.3-mm mesh net to promote water and sound exchange while preventing escape. The experiment took place in four black rectangular plastic tanks filled with 90 L (75 x 60 x 20 cm) of aged tap water at 20-22°C under a 12:12 light:dark cycle and equipped with an underwater loudspeaker (UW30 Electro Voice®) in the middle just below the water surface. Each tank received 18 glass microcosms placed on the bottom, equidistant from each other, and all at 20 cm from the speaker. We broadcast silence in two tanks (control) and broadband noise in the two others (noise treatment). Control *D. magna* therefore experienced laboratory background noise (see below for further detail). To account for the parental and genetic effects, half of the newborns from each single mother were allocated to the control and the other half to the treatment. They were fed every 2 days with 2 ml of algae (1g/L) directly in the glass microcosms with Pasteur pipette, and the water was changed once a week. Every day, the survival of each individual was checked and the offspring (when there were) were counted and removed (Fig. 1b). In the case of death during the eight first days (i.e., before the first hatching of the experiment), dead *D. magma* were replaced by 24h-newborns to keep the number of replicates. We used a total of 116 *D. magna* adults to obtain a total of 204 neonates, with 119 allocated to the control and 85 to the noise treatment (the different number of individuals between the two conditions resulted from different mortality rates in the eight-day individuals, i.e., the replacement previously described). Both in the control and the noise treatment, 39 individuals reached maturity and produced offspring at least once. The experiment lasted until the death of the last individual (the oldest *D. magna* survived 47 days).

**Figure 1.**
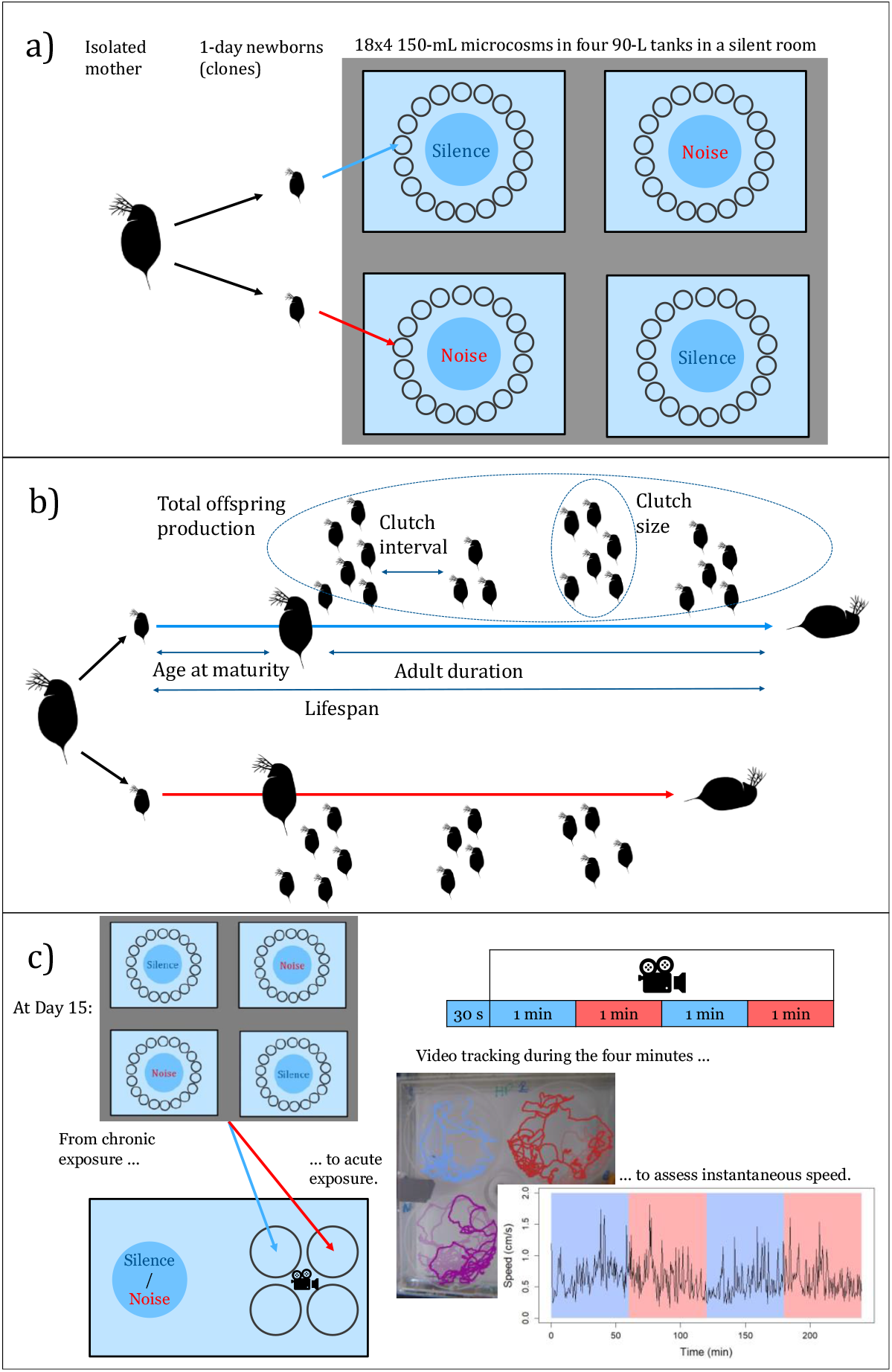
Setup. a) Experimental design of the experiment dedicated to *D. magna*’s fecundity and mortality. In the four tanks blue circles are loudspeakers, and the dark small circle represents microcosms closed with a net. b) Summary of all the measures made from birth to death on clonal individuals in the two treatments. c) Experimental design of the experiment on *D. magna*’s speed. Adapted from Prosnier et al. (2023).

From these daily and individual survival and fecundity data, we also analysed the effect of noise on populational growth using the Euler-Lotka equation, Σ*f*_*x*_*m*_*x*_*e*^−*rx*^= 1, with *f*_*x*_ the fecundity at age *x* (i.e., the number of offspring), *m*_*x*_ the survival at age *x* (i.e., the proportion of surviving individuals), and *r* the intrinsic rate of increase (Euler, 1760; Lotka, 1907; Corte s, 2016). Following this equation, three parameters were determined: the reproductive output *R*_*0*_ (*R*_0_ = Σ*f*_*x*_*m*_*x*_), the generation time 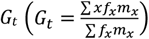, and the intrinsic rate of increase 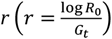 (Leung et al., 2007; Starke et al., 2021). Here, we calculated these three parameters by considering each treatment as a single population (Prosnier, Rojas, et al., 2023).

### Speed and size

We measured the speed and size of *D. magna* around day 15 (Fig. 1c). We randomly selected four individuals both from control and treatment and placed these four individuals in a 4-well dish (6 cm diameter x1.5 cm height). The dish was then positioned in a 20-L aquarium (20×25×40cm) filled with aged tap water, and exposed to natural light through a window. Using a dish allowed us to measure speed in two dimensions (horizontal plane). An underwater loudspeaker UW30 was located on the left and the dish was placed on the right side, on a brick to be in the middle of the aquarium. After an acclimatisation period of 30 s, a 4-min audio track (WAV file) was played, containing successively 1 min of silence, 1 min of noise, 1 min of silence and 1 min of noise.

*D. magna* were filmed using a GoPro Hero 4 Session camera and the videos were analysed using the Kinovea software (0.9.1 beta), which provided instantaneous speed every 33 ms. We used the videos to assess body size from the top of the head to the base of the caudal spine.

### Sound conditions

Two stereo WAV files were created for the experiment: one made of silence (to take into account the potential effects of the electromagnetic field) and the second of continuous broadband noise (100-20,000 Hz) generated in the Adobe Audition software (13.0.0.519, Adobe Systems Inc., Mountain View, CA, USA). The audio tracks were played by an UW30 underwater loudspeaker (Electro Voice®) connected to an amplifier (DynaVox® CS-PA 1MK), itself connected to a ZOOM H4next Handy recorder. To assess the acoustic spectra and sound levels in each glass-microcosm, we recorded the playbacks with a hydrophone (Aquarian Audio H2A-HLR Hydrophone, frequency response from 10 to 100 kHz) coupled to a Zoom® H4n, which was previously calibrated with a hydrophone (8104, Bru el & Kjær, Naerum, Denmark; sensitivity –205 dB Re 1 V μPa−1; frequency response from 0.1 Hz to 180 kHz) connected to a sound level meter (Brue l & Kjaer 2238 Mediator, Naerum, Denmark). Because the characteristics of the experimental units induce acoustic distortion, we adjusted the spectrum of the broadband noise (Fig. 2a) to make it as close as possible to a white noise between 100 and 20,000 Hz, using the equalizer of Adobe Audition 2020 software (13.0.0.519, Adobe Systems Inc., Mountain View, CA, USA). The mean Sound Pressure Level, SPL (in equivalent continuous sound level, Leq), within the crustacean earing range, 0.1-3 kHz (Duarte et al., 2021; Sole et al., 2023), was 90 dB RMS Re 1µPa^2^ in the control and 126 dB RMS Re 1µPa^2^ in the noise treatment (Fig. 2b), which is close to what has been reported in real freshwater systems with and without anthropogenic noise (Wysocki et al., 2007; Amoser & Ladich, 2010; Putland & Mensinger, 2020; Valenzisi et al., 2024). Note that the mean SPL over the 0-20 kHz frequency band is, respectively, around 118 and 128 dB RMS in the control and the treatment due to the energy in the low frequencies (0-100 Hz). To promote result comparison with other studies, we also estimated the dose of noise received by the daphnia over a 24-h period through the calculation of the Sound Exposure Level (SEL) (Slabbekoorn et al., 2010; Martin et al., 2019). The mean 24h-SEL for the broadband noise is 178 dB SEL_24h_ Re 1 μPa^2^.s for the control (149 dB SEL_24h_ for 0.1-3 kHz), and 188 dB SEL_24h_ 1µPa^2^.s for the treatment (186 dB SEL_24h_ for 0.1-3 kHz). Regarding the mobility experiment, another spectral correction was necessary using the method described above, as the experimental unit was different. The SPL values measured next to the 4-well dish in the 20-L aquarium were close to those obtained in the fitness experiment (93 dB RMS and 130 dB RMS Re 1µPa^2^ over the 0.1-3 kHz frequency band in the absence and presence of noise, respectively). For information, the 0.1-3 kHz SEL_1min_ was 115 dB Re 1µPa^2^.s for the control and 158 dB Re 1µPa^2^.s for the treatment. Due to the experimental setup, we were not able to measure, with an accelerometer, particle motion, although it is the main component of a sound for non-hearing species (Nedelec et al., 2016). However, relying on SPL to assess the difference in sound level between the treatments remains qualitatively relevant (Jones et al., 2022; Olivier et al., 2023).

**Figure 2.**
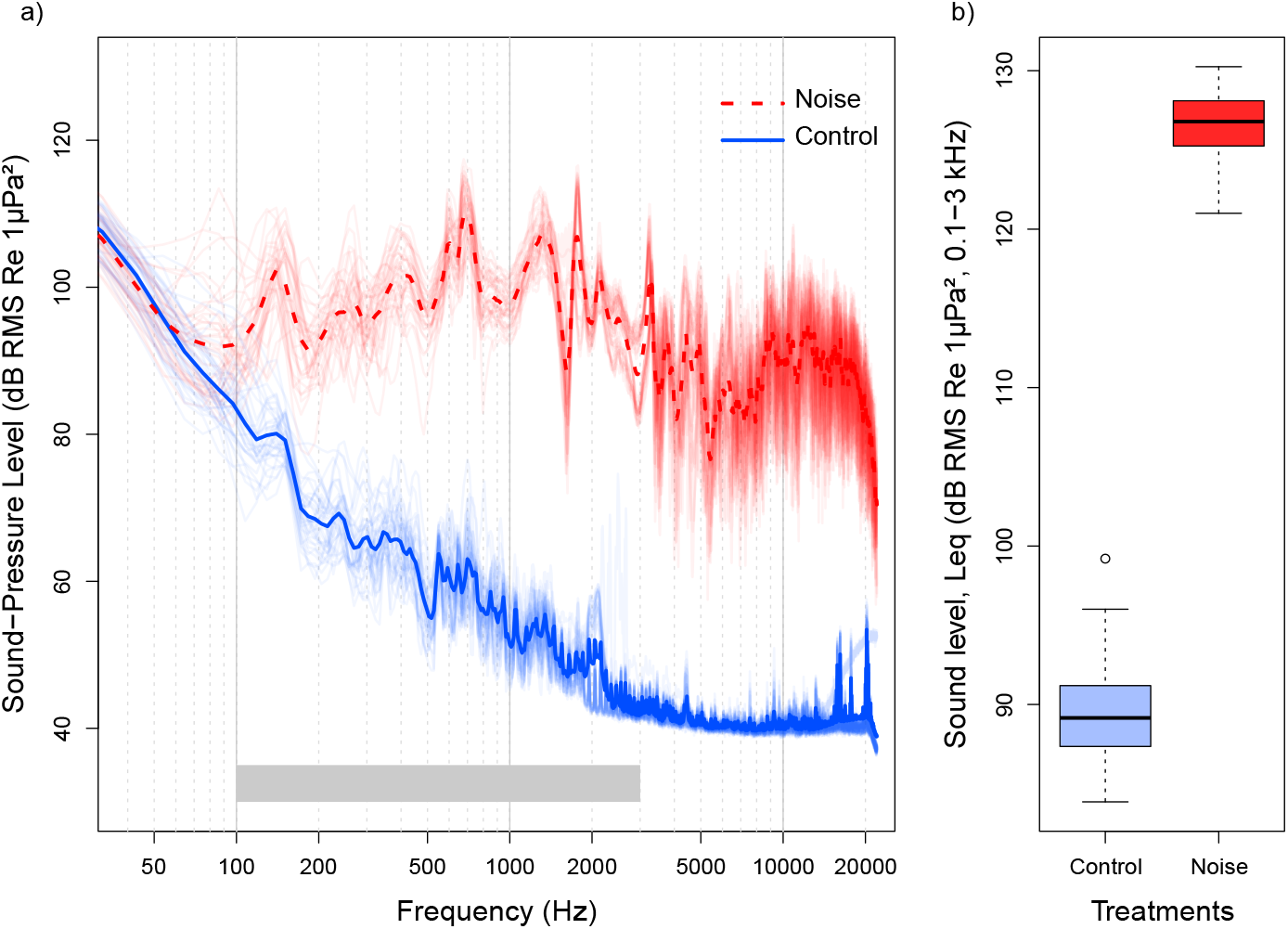
Acoustic conditions in the microcosms. a) Averaged (thick lines) sound pressure spectra for the control (full blue line) and the noise treatment (dashed red line). Transparent lines are the original sound spectra for each microcosm. The grey band indicates the crustacean earing range, between 0.1 and 3 kHz, according to Duarte et al., 2021. b) Box plot of the sound levels, within the crustacean earing range (0.1-3kHz) in all microcosms with central bars showing the medians and boxes the interquartile ranges. The dot is an outlier.

### Statistical analyses

Statistical analyses were performed using R software (version 4.2.2) with a significant threshold at 5%. According to our data, we separately analysed the effects of noise on mortality (death age, adult survival, and juvenile survival) and fecundity (age at maturity, clutch frequency, mean clutch size, and daily clutch size). We used the total production of offspring, considering both survival and fecundity, as a proxy for fitness. We performed a survival analysis (Log-Rank test) to compare survival (death age, adult duration, and survival of juveniles that never clutch) and age at maturity (first clutch age) between the two noise conditions. For the fecundity parameters, we considered only individuals that reached maturity. We used a linear mixed-effect model with clutch frequency (i.e., mean time between two clutches) and mean clutch size as response variables, the treatment as fixed effect, and tank and mother IDs (i.e., clone and maternal effect) as random effects. We analysed the effect of noise condition and age on the daily clutch size with a type II analysis of variance (ANOVA), followed by a Wilcoxon pairwise test between the two noise conditions for each age. The effect of the noise conditions on the total number of clutches and offspring along life were modelled with a generalized linear mixed-effects model, with tank and mother IDs as random effects and a log function as the link function for the Poisson distribution. We also estimated population growth in each noise condition using the Euler-Lotka equation (but with no statistical analysis due to the absence of population replicates). The difference in body size between the two noise treatments was tested with a t-test (after testing normality with a Shapiro test and homoscedasticity with a Bartlett test). For *D. magna*’s speed, we performed two analyses. First, we tested the effect of noise exposure (including the effects of size and age) on the average speed during the 4 min with a type II ANOVA (after controlling for normality and homoscedasticity). Then, we tested the effects of both chronic (i.e., with *D. magna* coming from the noisy microcosms) and acute (i.e., with *D. magna* coming from the noiseless microcosms) exposure to noise by comparing the mean speed during each 1 min period of treatment (from the 5^th^ to the 55^th^ second) using Tukey contrast on a linear mixed model, with individual ID as random effect.

## Results

### Fecundity and mortality

The survival of *Daphnia magna* was significantly higher with broadband noise compared to the control (p-value = 0.002, Fig. 3a, Table A1), as the adult duration was longer (p-value = 0.015) with four additional days (median value) with noise. There is no difference in the mortality of individuals dead before first clutch, i.e., the juvenile mortality (p-value = 0.88). Regarding fecundity, mean clutch size was significantly higher under noise than in the control (p-value = 0.005, Fig. 3b) but there was no significant difference in clutch frequency (p-value = 0.82, Fig. 3c) nor in age at maturity (p-value = 0.3), with 9 days for the two groups. Noise had no effect on daily clutch size (all p-values > 0.17, Fig. A1), which was only influenced by age (p-value < 0.001). Overall, noise exposure significantly increased the total number of clutches (p-value = 0.01) and the total offspring production (p-value = 0.009, Fig. 3d). Using the Euler-Lotka equation to study population growth confirmed that the reproductive output (*R*_*0*_) was higher for the noise treatment (39 offspring) compared to the control (12 offspring). The generation time (*G*_*t*_) seemed 3 days longer for the noise treatment (17 days) compared to control (13.9 days). The combination of these two parameters resulted in a higher intrinsic growth rate for the noise treatment (0.22 day^-1^) than for the control (0.18 day^-1^).

**Figure 3.**
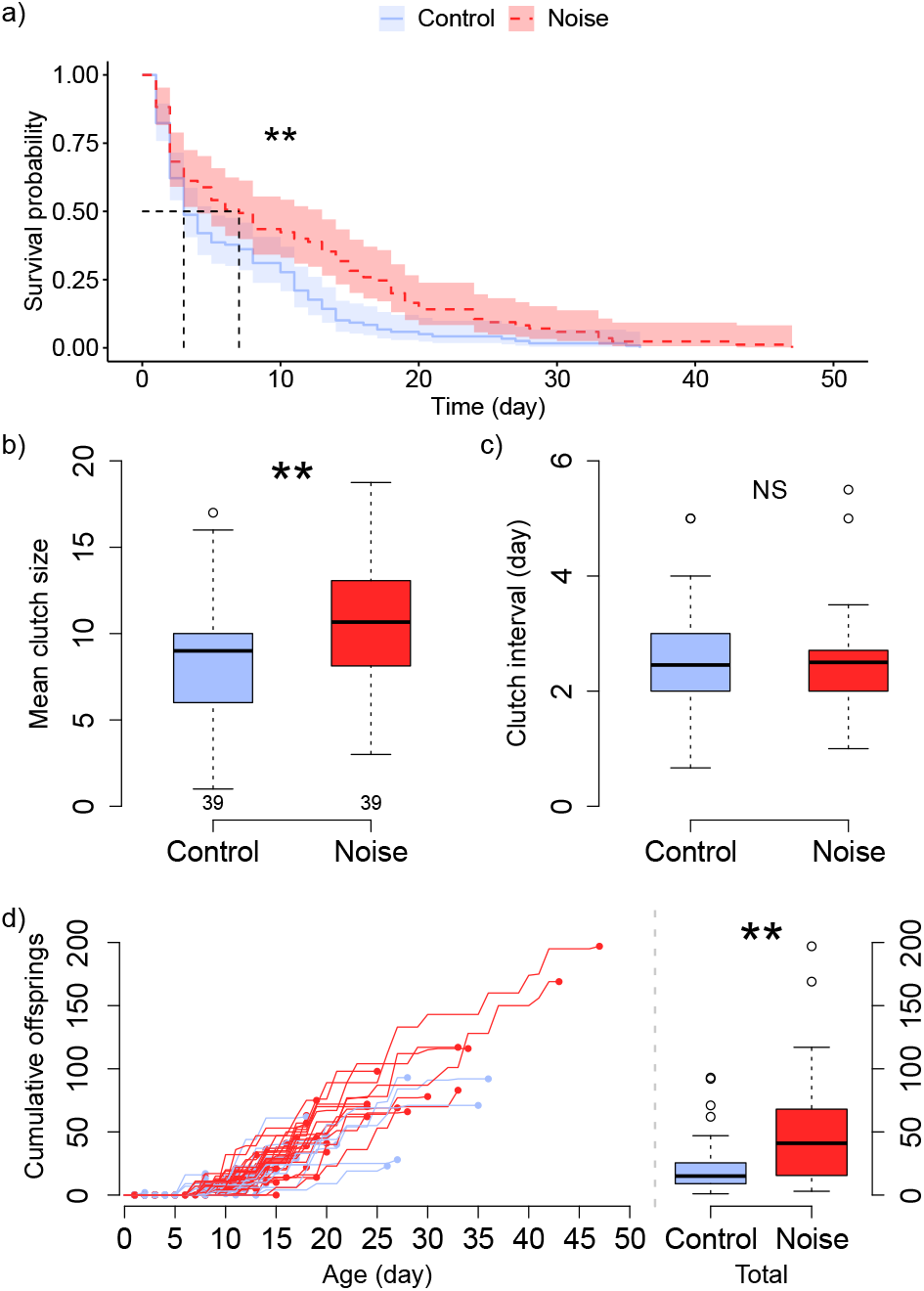
Survival and fecundity of laboratory-raised *Daphnia magna* exposed to broadband noise. a) Survival of *D. magna* according to the Kaplan-Meier method; b) mean clutch size; c) clutch frequency; d) dynamic of offspring production and total number of offspring during lifetime. Numbers in b) are the quantities of *D. magna* for the two treatments (light blue: control, dark red: broadband noise) for b-d. b-d) box plots with central bars for the medians, boxes for the interquartile ranges and dots for the outliers (> 1.5 times the interquartile range). Statistical analysis: ** *P* < 0.01; NS *P* > 0.1. See Table A1 for the statistical values.

### Size and speed

We did not observe any difference in body size between the two noise conditions (p-value = 0.675, Table A2). Regarding speed, *D. magna* chronically exposed to broadband noise in the microcosms were 16% slower than control ones, both in the whole experiment (p-value = 0.008, Fig. 4 and Table A3), and during the second period of silence (p-value = 0.031) – a result not explained by differences in size or age (p-values > 0.36). For the acute exposure (i.e., the 1-min of noise) there was no difference in mean speed between the two groups of *D. magna* in each acute exposure (p-values > 0.15), nor within the two groups of *D. magna* between each acute exposure (p-values > 0.88).

**Figure 4.**
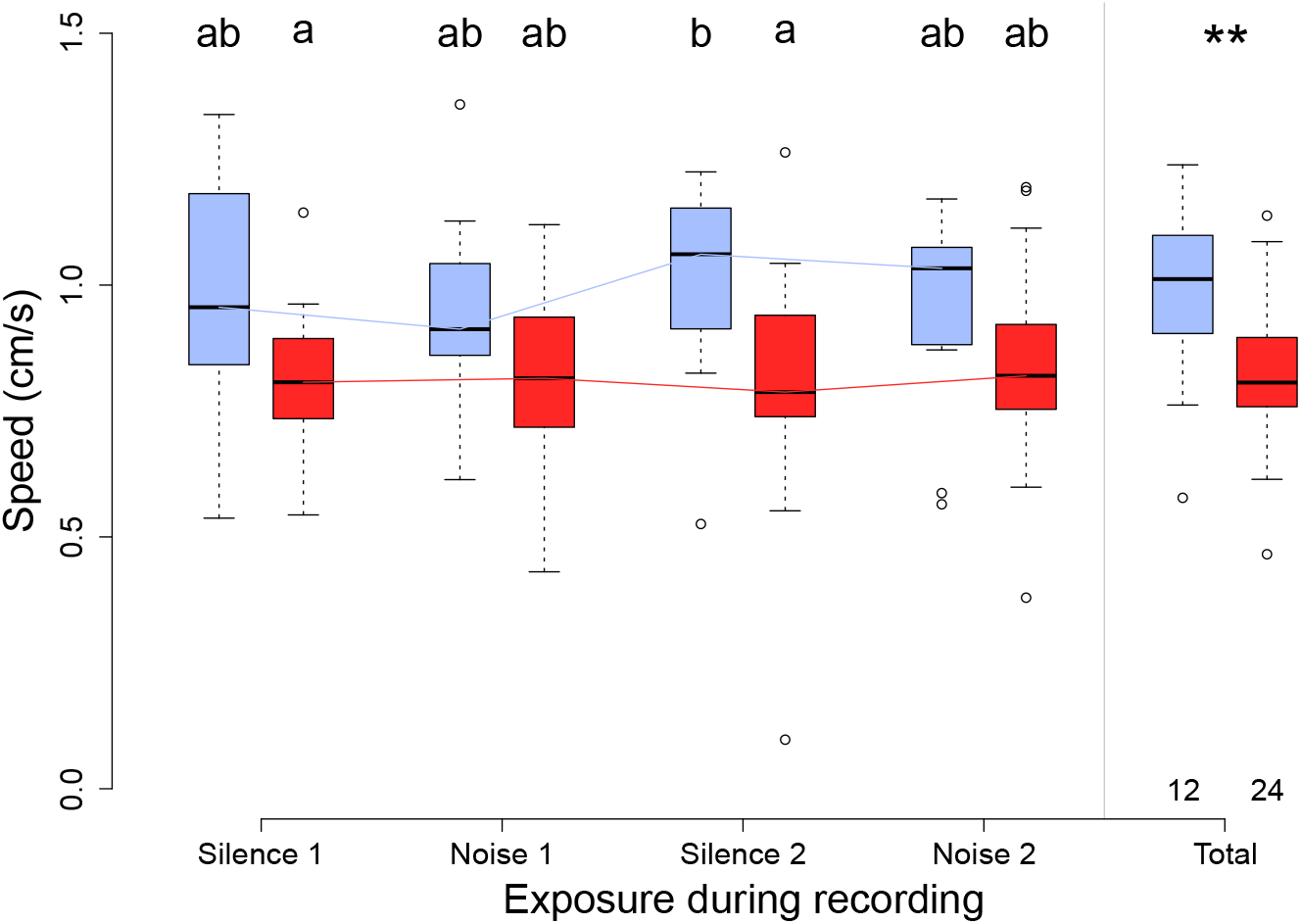
Swimming speed of *Daphnia magna* exposed to successive silent and noisy (broadband noise) periods and coming from control (in light blue) or noisy (in dark red) microcosms. Median and interquartile ranges (with dots for the outliers: > 1.5 times the interquartile range) of *D. magna*’s speed (in cm/s) during each of the four 1-min sequences of silence (Silence 1 and Silence 2) and noise (Noise 1 and Noise 2), and during the entire 4-min sequence on the right (Total). Numbers are the quantities of *D. magna* from each treatment. Paired data are shown with lines. The same letters indicate groups that are not significantly different at the level of 0.05 between the four sequences and the two groups of *D. magna*. ** *P* < 0.01. See Table A3 for the statistical values.

## Discussion

We did a playback experiment to study the effects of chronic continuous broadband noise on the fitness (fecundity and mortality) and mobility of laboratory-raised *Daphnia magna*, a widespread freshwater zooplankton coupling primary producers with higher trophic-level consumers of the pelagic food web. To our knowledge, it is the first study showing that long-term noise exposure could affect the fitness and behaviour of a zooplanktonic species (Prosnier, 2024). However, contrary to our prediction that constant stimulation would have long-term costs ultimately decreasing fitness, we found that exposed individuals had a higher survival and a larger clutch size, resulting in higher offspring production (i.e., a higher individual fitness). Noise also made *D. magna* less mobile, and it might be that the energy saved from reduced mobility, in laboratory conditions, was reallocated to survival and reproduction.

Previous laboratory investigations on *D. magna*’s response to anthropogenic noise did not provide evidence for any alteration. Sabet et al. (2015) used 6-min playbacks of continuous or intermittent 300-1500 Hz band-pass filtered white noise with sound pressure levels (SPL) elevated from 95 to 122 dB Re 1 μPa between the control and the sound treatments. Mobility was not affected despite a slight, but not significant, decrease in swimming speed under continuous noise, which could be consistent with our result. *D. magna* also seems insensitive to motorboat noise according to Rojas et al. (2023) where the authors did not find any change in swimming speed in response to 24 motorboat sounds distributed over a 45-min playback track with SPL values from 4.81 to 27 dB RMS Re 1 μPa above a 95-dB background noise. Using the same protocol as in the present study, Prosnier, Rojas, et al. (2023) assessed the fitness of laboratory-raised *D. magna* exposed from birth to death to a playback mimicking realistic boat traffic with various motorboats passing between 9 a.m. and 6 p.m., for a 2-h daily cumulative noise, with SPL values between 108 and 136 dB RMS Re 1 μPa^2^. They found no effect on the survival, clutch size, or offspring production. Our calculation of the dose of noise (estimated through SEL_24h_) received by the daphnia was quite equivalent between Prosnier, Rojas, et al. (2023) and the present study (185 and 186 dB SEL_24h_ Re 1 μPa^2^.s over the 0.1 - 3 kHz frequency range, for respectively the boat noise and the broadband noise; and 146 and 149 dB SEL_24h_ for the controls). The difference in the results between the two studies probably did not result from different doses of noise, but could be related to the respective temporal features of the two types of noise (continuous here and chronic intermittent in Prosnier, Rojas, et al. (2023)), which is not captured by SEL values. This raises the question of whether a continuous noise is more impactful than an intermittent one, contrary to what has been reported with some vertebrates (Nichols et al., 2015). A way to understand impact of sound exposure would be to conduct additional studies on *D. magna*’s mechanoreceptors. Although many studies have focused on copepods’ mechanoreceptors and responses to acoustic stimuli (Yen et al., 1992; Gassie et al., 1993; Lenz et al., 1996; Weatherby & Lenz, 2000; Buskey et al., 2002), this is not the case for cladocerans. Actually, only Klann & Stollewerk (2017) showed mechanoreceptors on the setae of *D. magna*. Given the importance of *Daphnia* as a biological model, it is surprising that there are not more studies confirming (or invalidating) their presence, localizations, structure, and functions. The present study shows that noise affects cladocerans and highlights the need to study their mechanoreceptors more closely, and then to experiment the dose-response relationship relative to noise pollution (Tyack & Thomas, 2019; Prosnier, 2024) to identify potential thresholds above which individual responses are triggered, or at least detectable.

In addition to the observed increased fitness, our experiment also revealed reduction of mobility, with both chronic and acute exposition. Chronic noise exposure in the microcosms resulted in reduced mobility that persisted during the two 1 min periods without noise. Acute noise exposure (i.e., only during the two 1-min periods of noise during the mobility experiment) caused a reduced mobility of the control individuals (similar to the speed of exposed daphnia). The distinct outcome observed during the first silence period may be attributed to an insufficient acclimatation period, which led to higher variability in the speed of the control individuals. Mobility is known to be linked with body size in the closely related *D. pulex* (Dodson & Ramcharan, 1991), which could potentially explain the chronic effect if chronic noise affects their development, but body size did not differ between the two noise conditions in the present study. It might be that continuous stimulation of the mechanoreceptors made *D. magna* less able to perceive their surroundings, resulting in lower mobility. A reduced mobility has already been reported for copepods exposed to an acute pressure drop (Vereide et al., 2024), and after a chronic exposure to a low-frequency noise (Tremblay et al., 2025). This is, however, the first documented behavioural effect of sound on *Daphnia* (Sabet et al., 2015, 2019; Rojas, Prosnier, et al., 2023), although Yuan et al. (2021) reported a reduction in speed following brief vibrational stimulation. Our findings raise questions about the role of sound level, as well as its temporal and spectral characteristics. For instance, the difference in fitness outcomes, between our results and Prosnier, Rojas, et al. (2023) should be due to the difference between our continuous noise and their boat noise. Although we did not assess food intake, *D. magna* were fed *ad libitum* so reduced mobility probably did not result in reduced energy input, as shown with copepods exposed to boat noise (174 dB SEL Re 1 µPa^2^) (Ku hn et al., 2023), and as it could be in natural populations (Read et al., 2014). Consequently, if we consider that energy input was preserved, then the energy saved from mobility might have been reallocated to tissue maintenance and reproduction, explaining the higher fitness. A trade-off between mobility and fitness was already reported for *D. magna* (Prosnier, Loeuille, et al., 2023) and for the codling moth *Cydia pomonella* (Gu et al., 2006). Further experiments should directly measure food intake, as done for copepods by Ku hn et al. (2023). Interestingly, the mechanoreceptors identified by Klann & Stollewerk (2017) are located on swimming and filtering setae, which are likely related to both mobility and feeding. Other important energetical parameters like growth rate, faeces production, oxygen consumption and energy content could be measured to study how noise influences energy allocation, for instance through the parametrization of a dynamic energy budget model (Nisbet et al., 2010; Sousa et al., 2010; Baas et al., 2018; Sherborne & Galic, 2020).

Evolutionary theory predicts investment in short-term reproduction when expectation of future offspring production decreases and under the perception of increased mortality risk (Williams, 1966). Such a life-history strategy, referred to as the *terminal investment hypothesis* (Clutton-Brock, 1984), might explain the noise-induced increase in fitness that appears counterintuitive at first sight. Fecundity compensation through early reproduction has already been reported in *D. magna* following exposure to the horizontally-transmitted microsporidian *Glugoides intestinalis* (Chadwick & Little, 2005) and to the sterilizing bacterial pathogen *Pasteuria ramosa* (Vale & Little, 2012). However, increased short-term reproduction is predicted to go with increased mortality under the terminal investment hypothesis and our study does not provide evidence for any mortality following exposure to noise.

The implications of our results for both field and laboratory studies should be explored in future research. First, it appears crucial to establish the dose-response relationships (Tyack & Thomas, 2019; Prosnier, 2024) with the key challenge of identifying critical thresholds for observed effects (fitness, behaviour, feeding, and energy…) while taking into account not only sound level (ideally in terms of particle motion, using average SPL and SEL) but also frequency (i.e., Power Spectral Density, PSD) and temporal features (i.e., continuous vs intermittent). Second, whether our results can be extrapolated to wild *D. magna* populations needs further investigation (Kok et al., 2023). Animals can show deviations in behaviour between the laboratory and the field (Tiselius, 1995; Prosnier, Loeuille, et al., 2023), and reduced mobility might have detrimental outcomes in the field as mobility is important to find food, to escape predators and more generally to do diel vertical migrations (Dodson et al., 1997; Larsson & Kleiven, 1997; O’Keefe et al., 1998; Roozen & Lu rling, 2001; Chang & Hanazato, 2003). Moreover, although we were only able to measure horizontal speed, many other behavioural aspects that are ecologically important and common in ecotoxicology could be investigated in future studies (Bownik, 2017; Bownik & Wlodkowic, 2021) – note that we attempted to perform experiments on vertical migration using a 3-m column, but noise calibration was difficult due to sound reflection (unpublished data). Knowing the indirect effects of this mobility reduction would be useful to understand how noise influences the fitness of natural population but also the entire zooplanktonic community through alterations in nested ecological interactions (Rojas, Desjonque res, et al., 2023; Rojas, Gouret, et al., 2023). Continuous noise can occur in the vicinity of certain infrastructures such as water pumping systems of compressors on the bank or wind turbines (Tougaard et al., 2020) and an interesting perspective to our work would be to study how water fleas use these noisy areas. Thirdly, water fleas represent key model species for ecotoxicological studies, as well as in many ecological and evolutionary studies (Ebert, 2005, 2022; Bownik, 2020; Abdullahi et al., 2022). Our results suggest that the background noise of rearing facilities matters (Davidson et al., 2007), in that it could interact with tested stressors – previous studies showed the interaction between chemical pollutants or temperature and acute or chronic noise on molluscs and arthropods (Shi et al., 2019; Stenton et al., 2022; Tremblay et al., 2025) –, influence population dynamics and the biological responses investigated. For instance, the background noise of rearing tanks affects both the physiology and the behaviour of the brown shrimp *Crangon crangon* (Lagarde re, 1982). Here, we observed that the cooling system of the experimental room induced an increase of around 10 dB inside the microcosm and the sound level of the control condition (silence) differed between the fitness experiment and the mobility experiment. Consequently, noise should be considered during the experimental setup, as other classical parameters like food and temperature, to avoid confounding factors (Harris et al., 2014).

## Acknowledgments

The authors would like to thank all those who contributed to the success of this experiment: Nicolas Boyer and Aure lie Pradeau for rearing the *Daphnia* and providing material, Le o Papet and Joe l Attia for their acoustic expertise, Marilyn Beauchaud and Paulo Fonseca for the acoustic calibration, and The ophile Turco and all the EYD (ENES Young Discussion) for the useful discussions. Authors also thank the five anonymous reviewers for their useful comments. The authors declare they had no funding for this research and were financially supported by their laboratory.

## Authors contributions

Conceptualization: All authors; Performing experiments: LP and OV; Analyses: LP, OV, and ER; Writing manuscript: LP, ER, and VM.

## Data, script, and code availability

Data, script, and code are available on Zenodo. DOI: 10.5281/zenodo.10614973 (Prosnier et al., 2024)

## Supporting information

Supplementary information is available after the references: Appendix: Table of statistics and supplementary figure

## Appendix

### Tables of statistics and supplementary figure

**Table A1.** Statistical results of the effects of chronic broadband noise on the mortality and fecundity of *Daphnia magna* (Fig. 3)

|  | Mortality |  |  | Reproduction |  |  | Fitness |  |
| --- | --- | --- | --- | --- | --- | --- | --- | --- |
|  | Survival | Juvenile mortality | Adult duration | Age at maturity | Clutch interval | Mean clutch size | Number of clutches | Number of offspring |
| df | 1 | 1 | 1 | 1 | 1 | 1 | 1 | 1 |
| $\chi^2/W$ | 9.9 | 0 | 5.9 | 1.1 | 0.0497 | 8.0469 | 5.9514 | 6.9065 |
| p-value | **0.002** | 0.88 | **0.015** | 0.3 | 0.82 | **0.005** | **0.01** | **0.009** |
| R<sup>2</sup> | 0.05 | 0 | 0.07 | 0.01 | NA | NA | 0.21 | 0.71 |

**Table A2.**
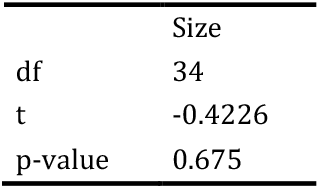
Statistical results of the effects of chronic broadband noise on the size of *Daphnia magna*.

|  | Size |
| --- | --- |
| df | 34 |
| t | -0.4226 |
| p-value | 0.675 |

**Table A3.** Statistical results of the effects of chronic broadband noise on the mobility of *Daphnia magna* (Fig. 4)

|  | Speed |
| --- | --- |
| df | 3-32 |
| F | 3.215 |
| p-value (trait) | **0.008** |
| p-value (size) | 0.361 |
| p-value (age) | 0.529 |
| R<sup>2</sup> | 0.232 |
| p-value (S1) | 0.152 |
| p-value (N1) | 0.351 |
| p-value (S2) | **0.031** |
| p-value (N2) | 0.689 |

**Figure A1.**
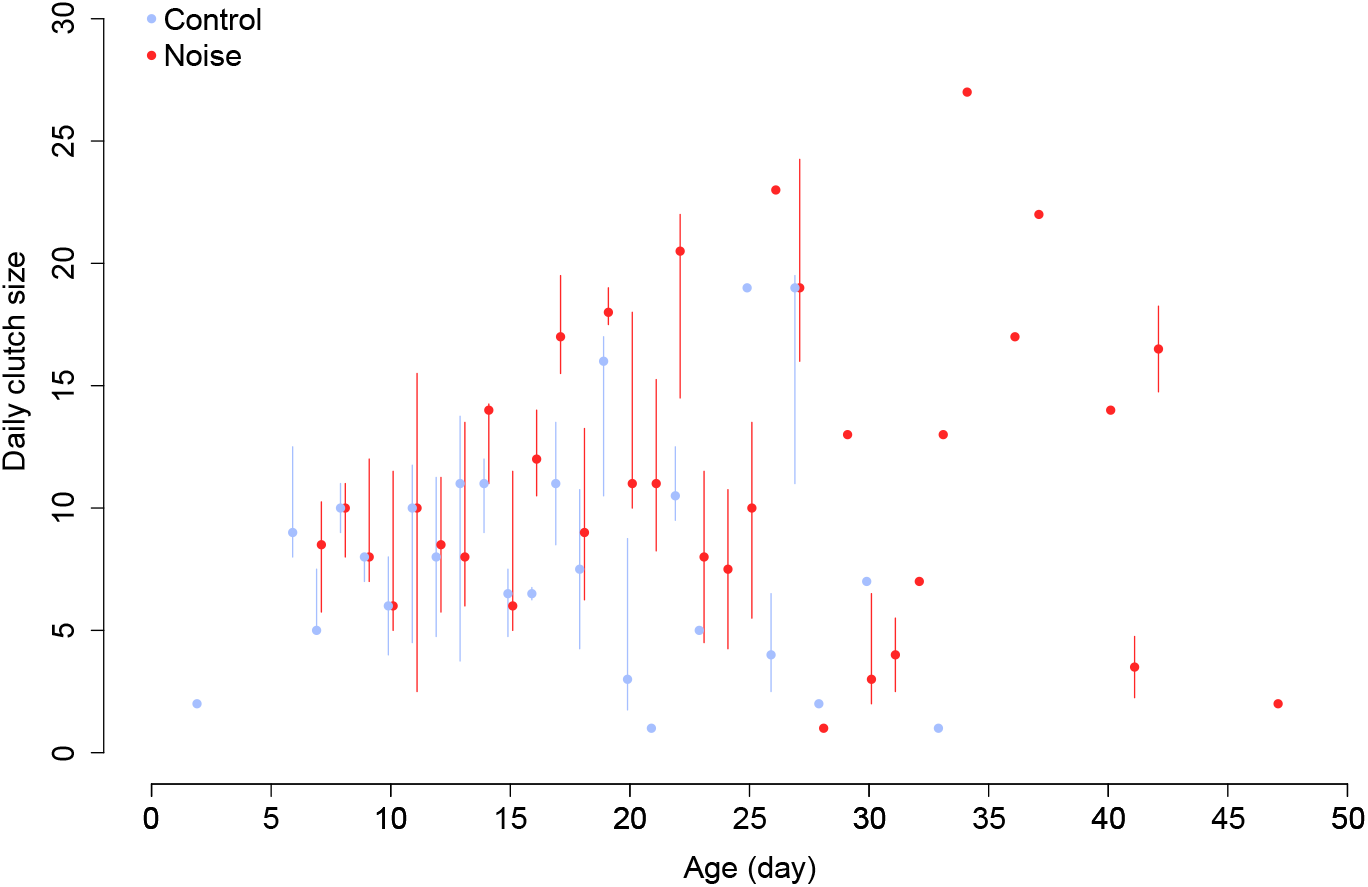
Effects of the noise conditions on *Daphnia magna*’s daily clutch size. Dots represent the median of the daily clutch size and lines are interquartile ranges. See Fig. 3a,d as complement for the number of individuals at each age. Note that there could be only one clutch for an age (i.e., no interquartile lines) or clutches only for one treatment (i.e., only one point for an age).

